# Stem cell-derived macrophages as a new platform for studying host-pathogen interactions in livestock

**DOI:** 10.1101/2021.09.10.459580

**Authors:** Stephen Meek, Tom Watson, Lel Eory, Gus McFarlane, Felicity J. Wynne, Stephen McCleary, Laura E.M. Dunn, Emily M. Charlton, Chloe Criag, Barbara Shih, Tim Regan, Ryan Taylor, Linda Sutherland, Anton Gossner, Cosmin Chintoan-Uta, Sarah Fletcher, Philippa M. Beard, Musa A. Hassan, Finn Grey, Jayne C. Hope, Mark P. Stevens, Monika Nowak-Imialek, Heiner Niemann, Pablo J. Ross, Christine Tait-Burkard, Sarah M. Brown, Lucas Lefevre, Gerard Thomson, Barry W. McColl, Alistair B. Lawrence, Alan L. Archibald, Falko Steinbach, Helen R. Crooke, Xuefei Gao, Pentao Liu, Tom Burdon

## Abstract

Infectious diseases of farmed and wild animals pose a recurrent threat to food security and human health. The macrophage, a key component of the innate immune system, is the first line of defence against many infectious agents and plays a major role in shaping the adaptive immune response. However, this phagocyte is a target and host for many pathogens. Understanding the molecular basis of interactions between macrophages and pathogens is therefore crucial for the development of effective strategies to combat important infectious diseases. We explored how pluripotent stem cells (PSCs) can provide a limitless *in vitro* supply of genetically and experimentally tractable macrophages from livestock. Porcine and bovine PSC-derived macrophages (PSCdMs) exhibited molecular and functional characteristics of *ex vivo* primary macrophages. Pig PSCdMs were productively infected by Porcine Reproductive and Respiratory Syndrome Virus (PRRSV) and African Swine Fever Virus (ASFV), two of the most economically important and devastating viruses in pig farming. Moreover, Pig PSCdMs were readily amenable to genetic modification by CRISPR/Cas9 gene editing applied in parental stem cells, or directly by lentiviral vector transduction. PSCs and differentiated derivatives therefore provide a useful and ethical experimental platform to investigate the genetic and molecular basis of host-pathogen interactions in livestock.

## Introduction

Recent global pandemics have focussed increased attention on the importance of understanding interactions between pathogens and their hosts. Pathogens carried by wild and farmed animal populations pose an evolving threat against both the primary hosts, and bystander species such as humans ^1^. The first line of defence against many pathogens is marshalled by macrophages, a phagocytic cell that operates as an essential arm of the innate immune system and regulator of the adaptive response ^2^. In some instances, however, macrophages are the preferred primary host cell targeted by pathogens, leading to dysregulation and disruption of the immune response ^3,4^. Indeed, infection and subversion of host macrophages is a common strategy used by viruses, bacteria and protozoans that compromise the health and productivity of the key livestock species, including pigs, cattle, sheep, and goats ^5–10^. The pathogens manipulate the host immune system to evade elimination by the host and thereby promote their survival and growth. A better understanding of interactions between host macrophages and pathogens is therefore critical for devising effective strategies to combat devastating, commercially important, diseases.

Domestic pigs are amongst the most numerous livestock species on our planet and under the conditions of contemporary farming management systems are susceptible to pathogen pandemics ^11^. In addition to the immediate economic and welfare impacts of disease outbreaks, infected herds can also serve as potential reservoirs for the development of candidate zoonoses ^12^. Pig macrophages in particular serve as targets and hosts for many important pathogens, including bacteria (e.g. *Salmonella enterica* serovars), protozoa (e.g. *Toxoplasma gondii*) and numerous viruses ^13,14^. Two key viruses, African Swine Fever Virus (ASFV) and Porcine Respiratory and Reproductive Syndrome Virus (PRRSV) target macrophages and are responsible for the most economically important infectious diseases in commercial pig farming ^4,15^. ASFV causes a lethal haemorrhagic fever, is highly contagious, and attempts to contain the most recent outbreak across Eastern Europe and Asia has resulted in the mass culling of millions of farmed pigs ^16,17^. PRRSV, in contrast, is endemic in most commercial herds and although not usually lethal causes reproductive failure and compromises productivity, imposing significant chronic welfare and economic costs ^15^. There are currently no commercially available vaccines or therapeutic interventions for ASFV and whilst live attenuated vaccines can be used to prevent severe disease from PRRSV, their efficacy is limited and linked to outbreaks caused by virulent revertants. How pathogens such as ASFV and PRRSV infect macrophages, deregulate their behaviour and control of the adaptive immune response, and ultimately destroy a key arm of the innate immune system, requires urgent attention. Access to physiologically relevant, tractable, experimental models with which to investigate these host-pathogen interactions is therefore vitally important.

Cell culture models play a crucial role in studying the molecular interactions between pathogens and their host macrophages ^18^. The current gold standard for these studies in pigs and ruminants are *ex vivo* macrophages harvested directly from slaughtered animals. These primary cultures exhibit very limited proliferation capacity and require constant replacement through a regular supply from donor animals, which incurs significant financial and animal welfare costs. Differences in the genetic background and immune status of animals, and inconsistencies in cell preparations also introduce significant batch-to-batch variability. Crucially, the primary cultures are not readily amenable to genetic modification, thus limiting prospects for rigorous functional dissection of macrophage-pathogen interactions. By contrast, transformed macrophages or heterologous cell lines are easier to genetically modify and can serve as surrogate hosts ^18,19^, but in many cases demonstrate only a subset of the key features of authentic macrophages.

Pluripotent stem cell (PSCs) lines are an alternative source of phenotypically “normal” macrophages ^20,21^. The stem cell lines are derived either directly from embryos or generated through induction of pluripotency through factor-directed reprogramming ^22^. PSCs can be expanded indefinitely in culture, are amenable to most gene manipulation techniques, and can be differentiated into a variety of cell types in culture including macrophages. The *in vitro*-derived macrophages closely resemble myeloid cells normally produced in the embryonic yolk sac that then colonise the foetus and contribute to the tissue resident macrophages of the adult ^23^. Despite their *in vitro* origins, PSC-derived macrophages exhibit similar phenotypic plasticity to *ex vivo* cells, and will adopt features of mature tissue resident characteristics in response to environmental cues ^21^. Mouse and human PSCs have been used previously to study the formation of the myeloid lineage, the role of macrophages as regulators of blood development, and innate immune responses to pathogens ^24–26^.

Until recently, translation of PSC-based technology to livestock species has been restricted by the difficulties in identifying culture conditions that robustly support PSC self-renewal. However, new studies have identified culture systems that support the derivation and propagation of embryo-derived PSCs from pig and cow ^27–29^. Here we describe the use of PSC lines as a source of macrophages and demonstrate their application as a culture model for studying host-pathogen interactions in the pig.

## Results

### PSC differentiation into macrophage-like cells

Pig PSCs were propagated on irradiated STO feeder support cells in pEPSC medium as described previously ^27^, and differentiated into macrophages using a 3 Phase protocol adapted from a method developed for human iPSCs (Figure 1A) ^30^. At the start of Phase 1, PSC cultures were pre-plated to remove residual feeder cells, aggregated by centrifugation, and then cultured for 4 days in suspension with bFGF, BMP4, VEGF and SCF to stimulate mesoderm differentiation and initiate formation of haematopoietic progenitors. In Phase 2, the day 4 aggregates were transferred to new culture dishes to allow attachment and cultured in IL-3 and CSF1 supplemented medium to promote expansion of haematopoietic progenitors and the development of macrophages. After 6 days of culture in Phase 2 (differentiation day 10), clusters of floating or loosely adherent cells containing vacuoles and short cell processes started to emerge from the cell monolayer. Between days 14-50, floating and loosely attached cells were collected every 3-5 days and re-plated in medium supplemented with CSF1 to promote the maturation of macrophages. Thereafter, medium containing batches of non-adherent cells were collected every 3-4 days for the next 4-5 weeks. The number of harvested cells peaked around day 28, and on average 7 × 10^6^ cells were produced from 10 aggregates, equating to 200 macrophages per input PSC.

**Figure 1.**
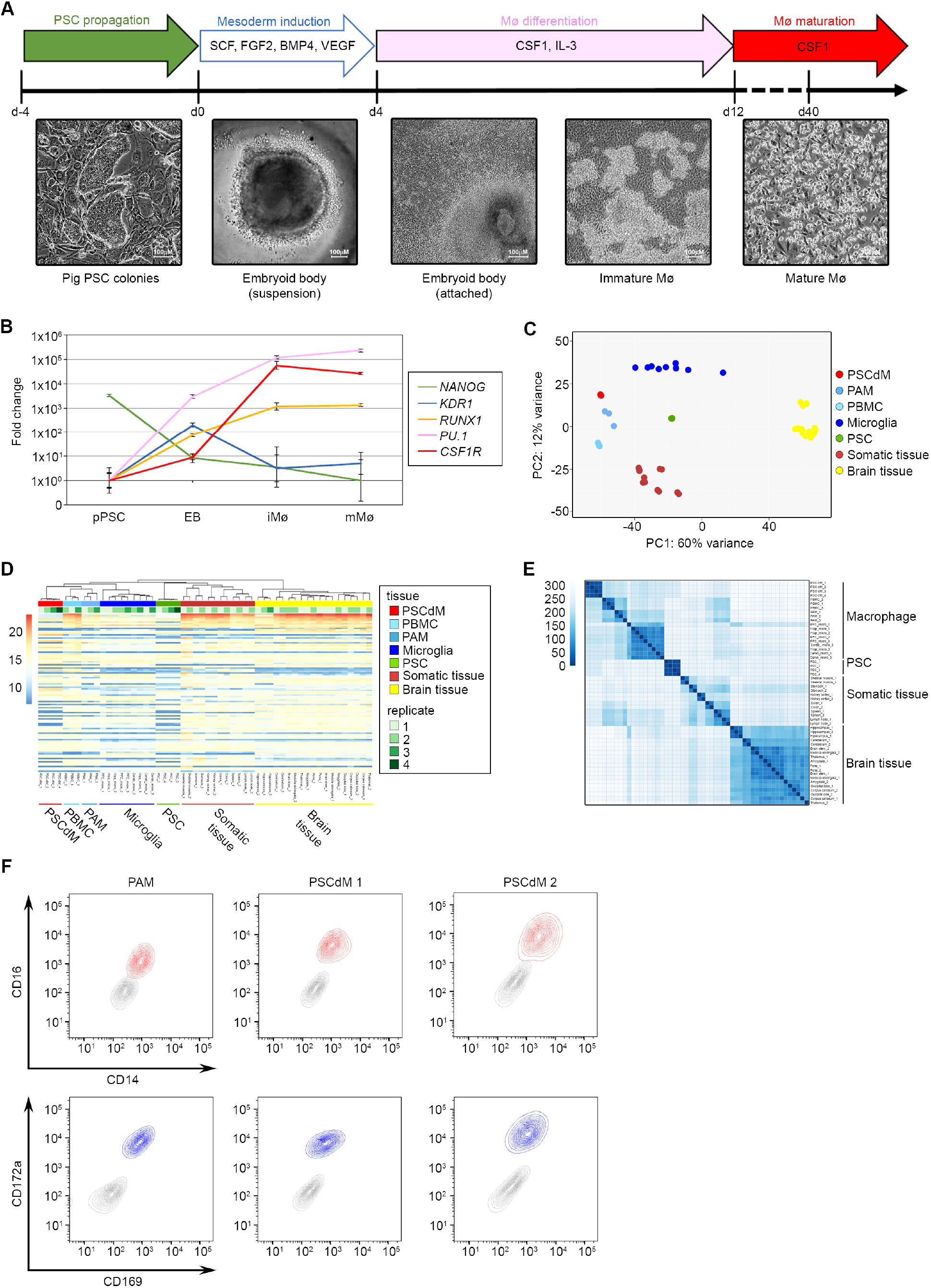
Generation and expression profiling of pig PSCdMs. **(A)** Schematic and timeline illustrating the differentiation protocol for deriving macrophages from pig PSCs. Cytokines used are indicated. Solid arrows indicate steps in which cells are attached: either on STO feeder cells (PSCs), gelatin (macrophage differentiation) or non-coated TC plastic (macrophage maturation). The hollow arrow representing mesoderm induction indicates that embryoid body formation was performed in suspension. Macrophages are first observed at day 9-12 and continue to be produced for approximately an additional 30 days, during which time clusters of floating immature macrophages can be collected every 3-5 days and matured in CSF1. Representative bright-field images are shown for the different cell morphologies generated at each stage. **(B)** RT-qPCR expression profile analysis of cells generated at each step of macrophage differentiation for markers of pluripotency (*NANOG*), early mesoderm induction (*KDR1*), HSC induction (*RUNX1*) and macrophages (*PU*.*1* and *CSF1R*). Mean and SD of two biological replicates from three experiments. **(C)** Score plot showing the first two principal components (PC1 and PC2) in tissue-specific gene expression. Based on PC1 and PC2 the data shows good separation of pig PSCs, *in vitro*-derived pig PSCdMs, *ex vivo*-derived pig alveolar macrophages (PAMs), microglia, brain and other tissue samples. **(D)** Heatmap of the hundred most highly expressed genes in pig tissues and cell lines. Lower expression levels are highlighted in blue and higher expression values in red. Biological replicates are indicated in green. Hierarchical clustering of the samples, shown as a tree at the top of the heatmap, was calculated using Euclidean distances between samples after transposing the variance stabilized expression data. **(E)** Heatmap showing the similarities between samples based on the Euclidean distances. The darker the colour, the closer the sample relationship is based on their expression profile. **(F)** Flow cytometry analysis comparing primary pig PAMs with *in vitro*-derived pig macrophages derived from two independent pig PSC lines (PSCdM 1 & 2) co-stained with CD14/CD16 (red) and CD169/CD172a (blue) relative to isotype controls (grey).

To track PSC fate transitions during this differentiation process, expression of genes characteristic of PSCs (*NANOG*), embryonic mesoderm (*KDR*), and haematopoietic progenitors (*RUNX1, PU*.*1*) and macrophages (*CSF1R*) were analysed in samples collected from starting PSC cultures and during Phases 1-3 (Figure 1B). Whereas transcripts of the stem cell marker *NANOG* declined immediately in Phase 1 indicating a rapid loss of pluripotency, expression of *KDR* increased and peaked at day 4, in line with the transient formation of embryonic mesodermal progenitors. By contrast, expression of haematopoietic/macrophage markers *PU*.*1, RUNX1* and *CSF1R* steadily increased through Phases 1 and 2, consistent with the formation of cells of the myeloid lineage. To assess the molecular phenotype of these macrophage-like cells produced in Phase 3 we compared their transcriptional profile against *ex vivo* pig macrophages and cells from a range of pig tissues by RNA sequencing. Multidimensional scaling provided a genome-wide overview of similarities between the different cell and tissue types, and demonstrated that the transcriptional profile of the *in vitro*-derived macrophages closely resembled that of *ex vivo* alveolar macrophages (PAMs) (Figure 1C). Hierarchical clustering analysis of the 100 most highly expressed genes further supported this conclusion (Figures 1D, E). The overall similarity between the transcriptional profile of the *in vitro* cells and *ex vivo* macrophages indicates that the pig stem cell-derived cells are closely related to endogenous macrophages.

Macrophages display a characteristic expression pattern of cell surface proteins. We examined expression of four typical macrophage proteins CD14, CD16, CD169 and CD172a, by staining Phase 3 PSC-derived cells with fluorophore conjugated antibodies recognising these cell surface proteins and analysing the cells by flow cytometry (Figure 1F) ^14^. The pattern of expression of all four proteins on differentiated cells generated from independent PSC lines was almost identical to that of *ex vivo* PAMs. Taken together with the transcriptome data, these results strongly indicate that the molecular profile of pig PSC derived cells (PSCdMs) is similar to *ex vivo* pig macrophages.

To assess the phagocytic activity of PSCdMs, these *in vitro*-derived cells and *ex vivo* PAMs were incubated with fluorophore labelled yeast particles (Phrodo beads) that fluoresce at a low pH, typically found in the acidic cell environment of the phagosome. Microscopic observation of Phrodo-treated PSCdMs showed that most of the cells had taken up fluorescent beads by 3 hr (Figure 2A). Quantitation of bead uptake during an 8hr incubation revealed that phagocytosis in PSCdMs was significantly higher than that observed in cultures of *ex vivo* PAMs (Figures 2B, C).

**Figure 2.**
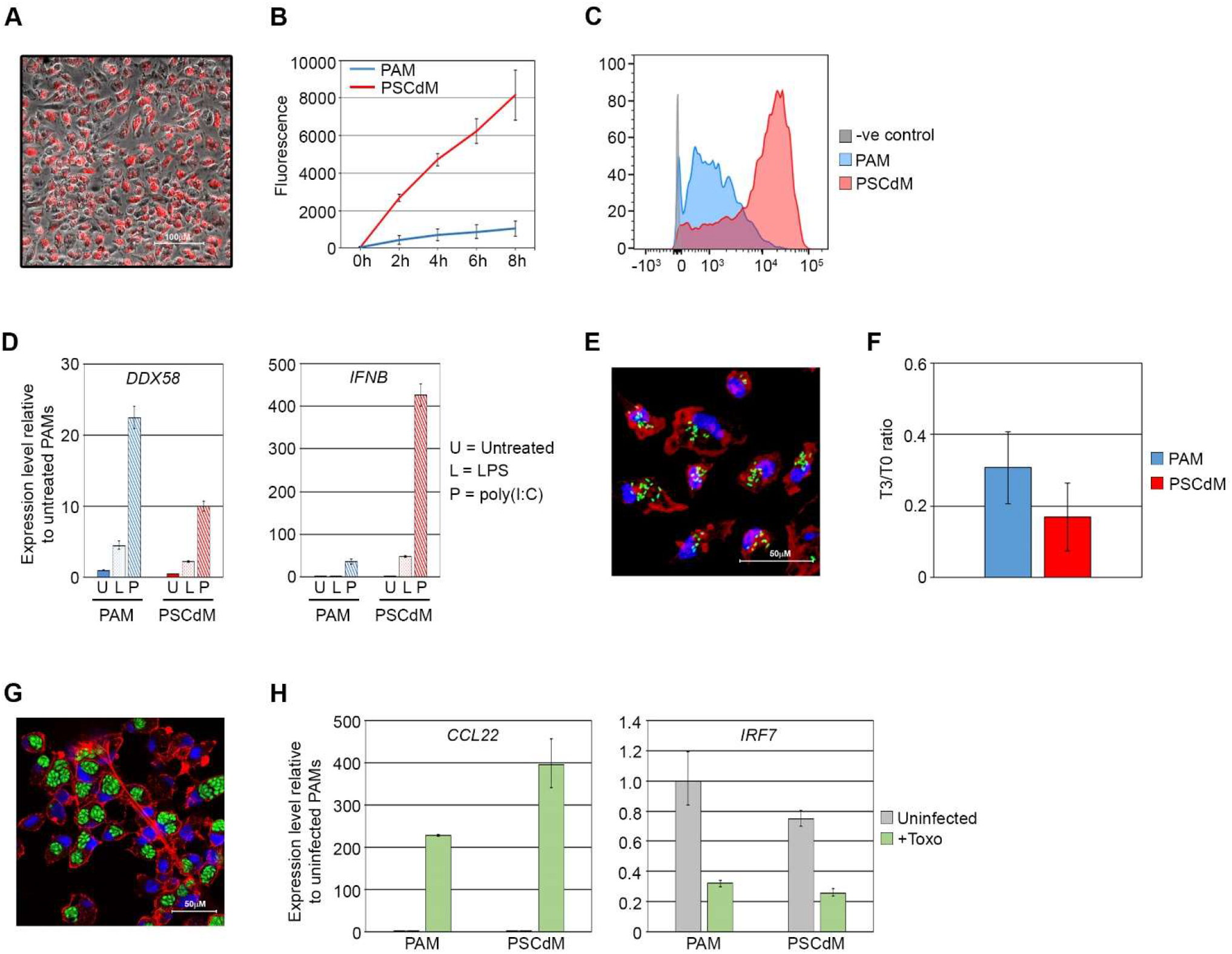
Functional validation of *in vitro*-derived pig PSCdMs. **(A)** Composite bright-field and fluorescent image showing phagocytosed pHrodo-Red beads fluorescing within pig PSCdMs. Image taken three hours after pHrodo bead addition. **(B)** Quantification of phagocytosis activity in pig PAMs (blue) and pig PSCdMs (red). Graph shows the level of pHrodo bead fluorescence between 0-8 h. Mean and SD of two pig PSCdM and one pig PAM line from three experiments. **(C)** Flow cytometry analysis of pig PAMs (blue) and PSCdMs (red) 8 h after pHrodo-Red bead addition relative to negative control cells (grey). **(D)** RT-qPCR analysis comparing *DDX58* and *IFNB* expression in pig PAMs and pig PSCdMs following 4 hrs pre-treatment with 200 ng/ml LPS or 25 μg/ml poly(I:C) relative to untreated controls. Mean and SD of three experimental replicates. **(E)** Confocal Z-stack projected image of pig PSCdMs 1 hr post-infection with EGFP-labelled *Salmonella typhimurium*. DNA is stained with DAPI (blue) and actin filaments with phalloidin (red). **(F)** Ratio of colony-forming *Salmonella typhimurium* recovered from infected pig PAMs and pig PSC-dMs at 3 hr post-infection relative to T0. Mean and SD of duplicate plates from two dilutions. **(G)** Confocal image of pig PSCdMs 24 hr post-infection with EGFP-labelled *Toxoplasma gondii*. DNA is stained with DAPI (blue) and actin filaments with phalloidin (red). **(H)** RT-qPCR analysis of *CCL22* and *IRF7* expression in uninfected and Toxoplasma gondii-infected pig PAMs and pig PSCdMs. Mean and SD of three experimental replicates.

To determine whether the PSCdM differentiation procedure could be applied to livestock PSCs more broadly, we tested the differentiation protocol on bovine PSCs (Figure S1A) ^28^. Bovine stem cells were propagated on mouse embryonic fibroblast feeders in N2B27-based medium containing FGF, Activin and the Wnt inhibitor IWR-1 ^31^. Upon differentiation using the PSCdM protocol, macrophage like cells emerged after 10-14 days in Phase 2 culture, within a similar time frame to pig PSCdMs. The bovine cells exhibited typical macrophage morphology, expressed the key macrophage markers *CSR1R, PU*.*1* and *RUNX1*, and were highly phagocytic, matching *ex vivo* primary bovine alveolar macrophages and indicating that PSCs could also serve as a useful source of bovine macrophages in future studies (Figures S1B, C).

### Response of pig PSCdMs to pathogens

To assess the potential of stem cell-derived macrophages for studying host-pathogen interactions, we challenged pig PSCdMs with biologically and economically relevant pig pathogens. To first determine whether PSCdMs recognise pathogen-associated molecular patterns (PAMPs), and initiate appropriate downstream immune responses we exposed the cells to either the synthetic dsRNA analog poly(I:C) or bacterial lipopolysaccharide (LPS) which are recognised by Toll-like receptors 3 and 4 respectively. The innate immune response gene *DDX58* (*RIG-1*) and the type I interferon gene *IFNB* were upregulated in PSCdMs treated with both poly(I:C) and LPS to a similar degree as *ex vivo* PAMs (Figure 2D) implying that PSCdMs would respond when challenged by bacterial and viral pathogens. To examine how PSCdMs react when challenged with live bacteria, and their capacity to resolve an infection, PSCdMs and control *ex vivo* PAMs were incubated with a *Salmonella enterica* serovar Typhimurium strain expressing enhanced green fluorescent protein (EGFP) ^32^. Internalisation of fluorescent *Salmonella* occurred within 1 hr after addition of the bacteria, and net intracellular survival, as assayed by a dilution gentamicin-protection assay, had declined to a similar degree in both PSCdMs and PAMs at 3 hrs, demonstrating an effective bactericidal capacity of the *in vitro* derived macrophages (Figures 2E, F). Comparable results were also obtained upon infection of PSCdMs and PAMs with *Escherichia coli* strain TOP10 (Figure S2).

The obligate intracellular parasite *Toxoplasma gondii* infects macrophages in a number of livestock species including pigs. It is also carried in most domestic cats and can cause severe disease when transmitted to immunocompromised humans ^13^. To test the response of PSCdMs to this pathogen, macrophages were incubated overnight with an EGFP-labelled strain of *Toxoplasma gondii* ^33^, and then examined by fluorescent microscopy (Figure 2G). Most PSCdMs contained identifiable rosettes of the EGFP-labelled protozoan, indicating that the macrophages were efficiently infected by *T. gondii*, and enabled replication of the parasite. *Toxoplasma* manipulates the host anti-microbial response by disrupting interferon signalling, and promoting an anti-inflammatory state ^34,35^. RT-qPCR analysis showed that in PSCdMs infected with *Toxoplasma* expression of *IRF7* (Interferon regulatory factor 7) was downregulated and expression of the anti-inflammatory chemokine *CCL22* was upregulated, mirroring the response obtained in PAMs and supporting the notion that the reaction of PSCdMs to colonisation by this intracellular parasite is comparable to *ex-vivo* porcine macrophages (Figure 2H).

African Swine Fever Virus (ASFV) and Porcine Reproductive and Respiratory Syndrome Virus (PRRSV) specifically infect pigs, primarily targeting the macrophage ^4,15^. PSCdMs were incubated with ASFV and at 24-48 hrs immunocytochemical detection of the p72 viral capsid protein and the formation of haemadsorption rosettes demonstrated that PSCdMs were readily infected by ASFV (Figures 3A, B). Quantitation of viral DNA in culture supernatants recovered from ASFV infected PSCdMs and *ex vivo* macrophages was performed by qPCR and showed that ASFV growth was efficiently supported by PSCdMs (Figure 3C). The productive infection of PSCdMs was confirmed using a TCID_50_ serial dilution assay, and together with the PCR result, established that the *in vitro*-derived macrophages can serve as effective hosts for propagating ASFV (Figure S3A).

**Figure 3.**
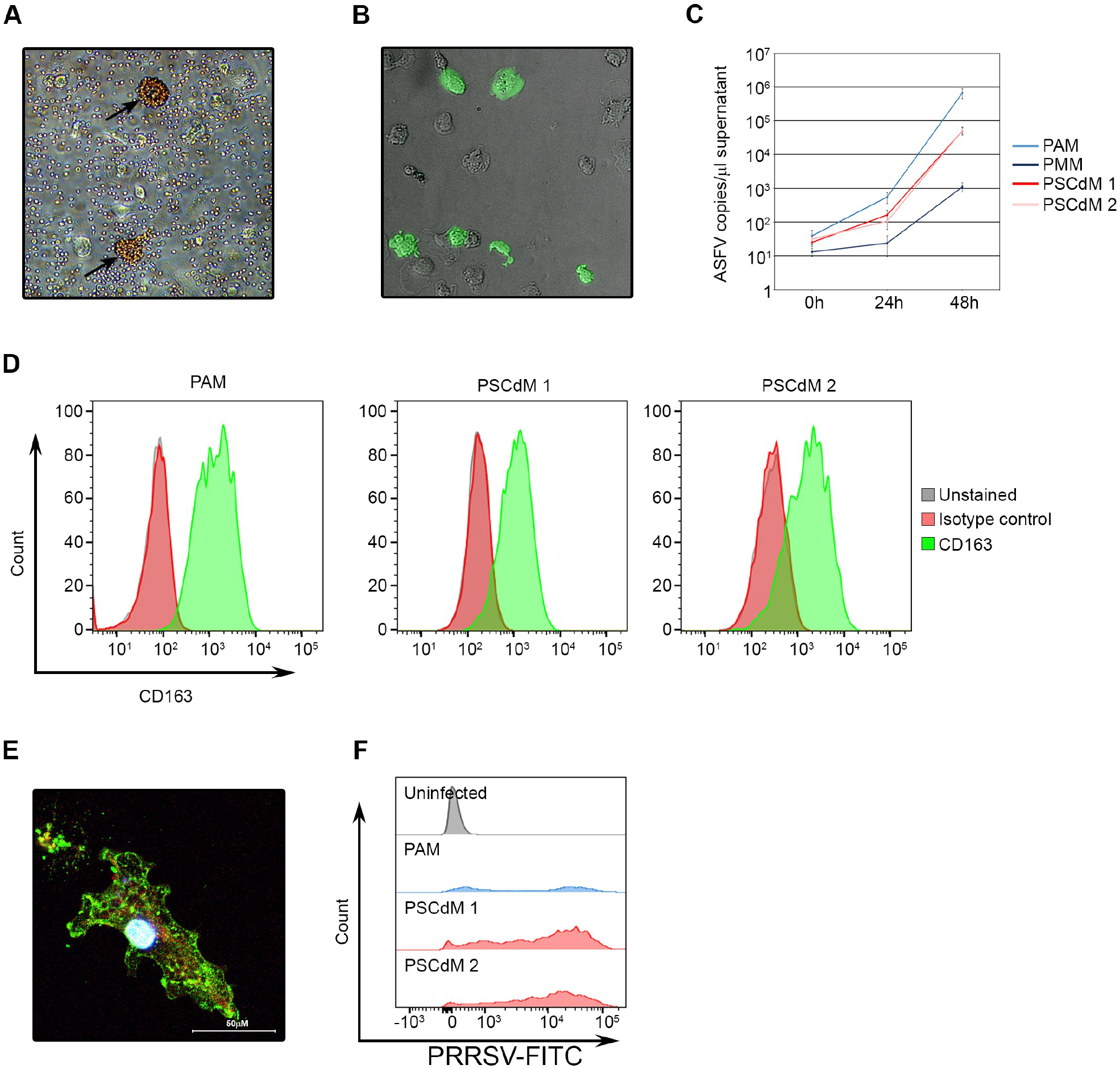
Viral infection of pig PSCdMs. **(A)** Bright-field image from a haemadsorption assay of ASFV-infected pig PSCdMs, 24 hr post-infection. Red blood cells can be seen aggregating around two pig PSCdMs (black arrows). **(B)** Composite bright-field and fluorescent image of ASFV-infected pig PSCdMs, 24 hr post-infection, stained for p72 viral protein (green). **(C)** RT-qPCR analysis of ASFV levels (genome copies) present in supernatants from pig PAMs and pig PSCdMs 24 h and 48 h post infection. Mean and SD of four experimental replicates. **(D)** Flow cytometry analysis of cell surface CD163 (green) on pig PAMs and pig PSCdMs derived from two independent PSC lines relative to isotype control (red) and unstained cells (grey). **(E)** Confocal image of a PRRSV-infected pig PSCdM, 19 h post-infection and stained for PRRSV nucleocapsid protein (green). DNA is stained with DAPI (blue) and actin filaments with phalloidin (red). **(F)** Flow cytometry analysis for PRRSV nucleocapsid protein in PRRSV-infected pig PAMs (blue) and two independent pig PSCdM lines (red) 18 h post-infection and relative to uninfected pig PSCdM (grey).

Efficient PRRSV infection of pig macrophages is mediated by the CD163 haemoglobin-haptoglobin scavenger receptor ^36^. Although *CD163* mRNA expression in PSCdMs as measured by RT-qPCR was ∼ 20% of that in PAMs, CD163 cell surface protein could be detected on the majority of PSCdMs by flow cytometry (Figure 3D, S3B). PSCdMs and PAMs were incubated with PRRSV (SU1-BEL) and infection was determined by measuring PRRSV p63 nuclear capsid protein expression by microscopy and flow cytometry (Figures 3E, F). PSCdMs were infected efficiently by PRRSV in line with their general pattern of CD163 protein expression. Between 18-24 hrs PSCdMs and PAMs began to lyse due to the cytopathic effects of PRRSV, and most macrophages were dead at 48-72 hrs. To confirm that PSCdMs support replication and production of infectious PRRSV, the culture supernatants were collected at intervals up to 72 hrs after infection and used to initiate a secondary infection on target PAMs. Flow cytometry analysis for PRRSV p63 expression in these secondary infections demonstrated that PRRSV production by PSCdMs, although not evident at 6 hrs, was readily detected at 24 hrs and at levels equal to or greater than produced by primary PAMs (Table 1). Collectively these results indicate that PSCdMs are infected by and respond to key pig pathogens, and can represent a useful experimental model to study host-pathogen interactions.

**Table 1.** Infection of PAMs with PRRSV cell supernatants harvested from PAMs and pig PSCdMs.

| Cell line supernatant | 6h | 24h | 48h | 30h | 72h |
| --- | --- | --- | --- | --- | --- |
| PAM | 0.12 | 43.6 | 43.8 | 46.6 | 36.2 |
| PSCdM 1 | 0.24 | 44.2 | 41.7 | 41.7 | 41.2 |
| PSCdM 2 | 0.36 | 49 | 54.6 | 47.9 | 48.0 |

Data represents the percentage of PRRSV nucleocapsid protein positive cells relative to uninfected controls.

### Genetic engineering of pig PSC-derived macrophages

A useful feature of established rodent and human PSC lines is that they are amenable to many contemporary genetic engineering techniques. To assess the feasibility of ribonucleotide protein (RNP) CRISPR/Cas9-mediated gene editing in pig PSCs we first targeted an EGFP-Puro knock-in reporter transgene into the non-essential pluripotency-associated *REX1* gene using a PITCh-based strategy ^37–39^ (Figure S4A). PSCs were electroporated with Cas9/gRNA RNP complexes that cut at the translation termination codon of pig *REX1* together with a PITCh vector directing integration of a 2A-EGFP-IRES-Puro cassette in tandem with the *REX1* open reading frame, allowing puromycin selection of correctly targeted cells. Puromycin resistant clones carrying the EGFP knock-in construct were isolated after 10-14 days selection and expanded to establish stable cell lines (Figure S4B). Cells within the undifferentiated *REX1-EGFP* PSC colonies expressed the *REX1-EGFP* reporter uniformly and, as predicted, downregulated its expression upon differentiation (Figures S4C, D).

We next deleted the pig gene encoding Interferon regulatory factor 3 (IRF3). This latent cytoplasmic transcription factor is activated in response to pathogens and plays a role in the induction of an interferon-mediated antiviral response ^40^. Disruption of *IRF3* would be predicted to increase viral replication, by uncoupling the endogenous antiviral response. To eliminate IRF3 from pig macrophages, PSCs were electroporated with a pair of Cas9/gRNAs RNP complexes designed to delete the entire *IRF3* coding region by cutting immediately after the ATG translation start codon and 3 bp after the termination codon (Figure 4A). PSC clones were isolated by limiting dilution cloning and screened by PCR for deletion of the *IRF3* exons (Figure 4B). On this basis, 46% of picked clones carried a deleted *IRF3* gene, and 4% were deleted on both alleles (Table 2). Three independent *IRF3* knock-out (KO) clones were expanded for further analysis, and all three differentiated to generate PSCdMs (Figure 4C). Analysis by qRT-PCR confirmed the loss of *IRF3* mRNA expression in KO macrophages (Figure 4D). To determine how the absence of IRF3 affects the response of pig PSCdMs to virus, the parental wild-type control and KO cell lines were incubated with PRRSV and after 24 hrs the amount of virus in cells was analysed by measuring p63 nuclear capsid protein expression by flow cytometry (Figures 4E, S5). PSCdMs were also pretreated with poly(I:C) prior to infection, to assess how IRF3 deficiency affects the induction of an antiviral state. Poly(I:C) binding to TLR3 mimics RNA virus infection and leads to activation of the IRF3 protein, which in turn increases transcription of *IFNB* and the induction of a protective antiviral state. PRRSV infection in the three untreated IRF3 KO cells was similar to the parental cell line, and pre-treatment with poly(I:C) reduced PRRSV infection dramatically in both types of PSCdMs, demonstrating that IRF3 was not essential for effecting an antiviral state in pig macrophages (Figures 4E, S5). However, the higher levels of virus detected in poly(I:C) treated KO cells indicated that establishment of the antiviral state was less effective in the absence of IRF3.

**Figure 4.**
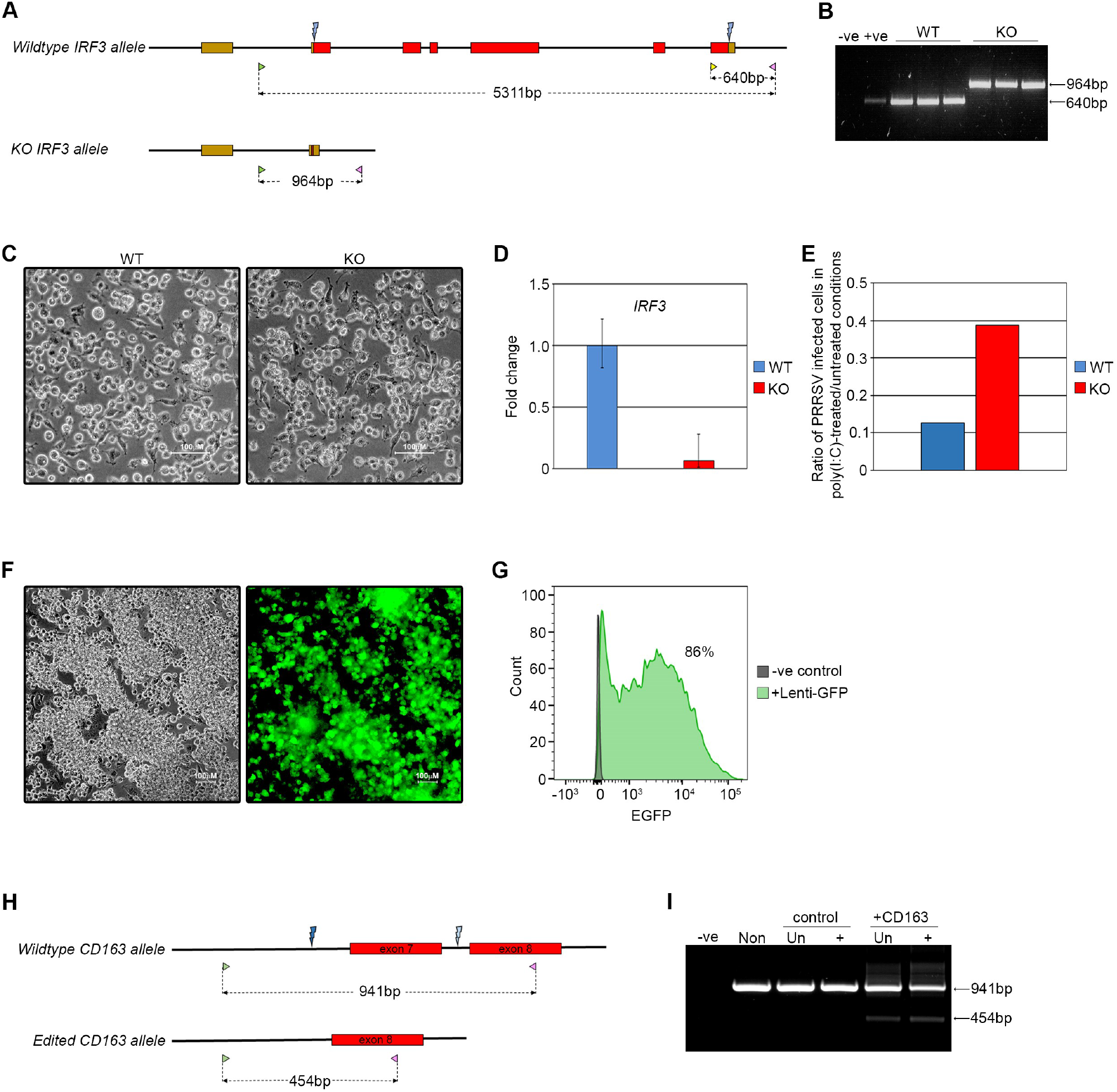
CRISPR-Cas9 editing in pig PSCs and PSCdMs. **(A)**CRISPR/Cas9 editing diagram showing wild-type (top) and knock-out (bottom) pig *IRF3* alleles. The entire *IRF3* coding sequence was deleted using a pair of guides (blue lightning bolts) designed to cut immediately after the initiation codon and 3 bp upstream of the stop codon. Coding exons are shown as red boxes, non-coding genomic sequence as thick black lines and 5’ and 3’ UTRs as brown boxes. Genotyping was performed using a pool of two forward primers (green and yellow arrowheads) and one reverse primer (pink arrowhead), where the yellow forward primer located in exon 6 is specific for the wild-type allele. Expected PCR product sizes are indicated. **(B)** Genotyping PCR result using the primer pool indicated in panel A. Three wild-type (WT) and three *IRF3* knock-out (KO) clones generated the expected product sizes (note that the additional 5311 bp wild-type product was never detected under these PCR conditions). Water was used as the negative control (-ve) and parental pig PSC genomic DNA as a positive control (+ve). **(C)** Bright-field images of pig PSCdM generated from wild-type (WT) and *IRF3* knock-out (KO) clones. **(D)** RT-qPCR analysis for IRF3 expression in pig PSCdMs from wild-type (WT) and IRF3 knock-out (KO) clones. Mean and SD of three biological replicates. Wild-type sample data is composed of macrophages derived from two PSC lines and one wild-type clone. **(E)** Ratio of PRRSV-infected WT and IRF3 KO pig PSCdMs in poly(I:C)-treated:untreated conditions. Macrophages were pre-treated with poly(I:C) (25 μg/ml) for 3 h prior to infection. Data represents ratio of parental line and mean ratio of three KO clones. **(F)** Brightfield and fluorescent images of lenti-EGFP-transduced pig PSCdMs, 7 d post-transduction. **(G)** Flow cytometry analysis of lenti-EGFP-transduced pig PSCdMs 7 d post-transduction relative to non-transduced pig PSCdMs. **(H)** CRISPR/Cas9 editing diagram showing wild-type (top) and edited (bottom) pig *CD163* alleles. Coding exons are shown as red boxes and non-coding genomic sequence as thick black lines. A pair of guides (blue lightning bolts) have previously been shown to delete exon7 ^55^ resulting in a 487 bp deletion that can be detected by a PCR screen using primers designed to flank the deleted region (green and pink arrowheads). Expected PCR product sizes are indicated. PCR analysis for lentiviral-mediated CD163 editing in pig PSCdMs. Macrophages were transduced with a Cas9-expressing lentivirus together with either a lentivirus expressing CD163 guide RNAs or an empty vector control. Transduced cells were isolated by FACS 7 days post-transduction for BFP expression from the guide RNA-expressing lentivirus. Unsorted and BFP^+ve^ macrophages were PCR amplified using the screening strategy in panel H. Water (-ve) and non-transduced cells (Non) were included as controls.

**Table 2.** *IRF3* editing efficiency.

| No. of colonies picked | Wild type | Heterozygous | Homozygous |
| --- | --- | --- | --- |
| 80 | 43 (54%) | 34 (42%) | 3 (4%) |

PSC culture provides opportunities for scalable production of normal and genetically modified macrophages suitable for larger genetic screens. To assess the feasibility of using pig PSCdMs in these type of experiments we first examined whether PSCdMs could be efficiently transduced by lentiviral vectors, since this virus-based system is commonly used for delivery of gRNA libraries^41^. Early Phase 3 PSCdMs were infected with a lentivirus expressing EGFP and flow cytometry of transduced cultures after 96 hrs showed that ≥ 95% of the surviving macrophages expressed high levels of EGFP (Figures 4F, G). We next-transduced PSCdMs with two lentiviruses, one directing expression of the Cas9 protein, and the other expressing the BFP fluorescent protein and two gRNAs designed to delete exon 7 of *CD163*. Five days after transduction with the lentiviruses, unsorted and BFP positive sorted cells purified by FACS were analysed by genomic PCR. Efficient deletion of *CD163* exon7 was detected in the transduced population and was enriched in the BFP positive fraction. Taken together these results demonstrate the use of pig PSCs and PSCdMs as a platform for interrogating gene function and developing genetic screens for investigating host-pathogen interactions in the pig (Figure 4H, I, S6).

## Discussion

The derivation of pluripotent stem cells from livestock species promises to advance prospects for studying the basic biology of these animals, and the development of strategies to improve their health and resistance to disease^42,43^. Livestock PSCs are a potentially limitless, and ethically unencumbered source of normal cells, and enable precision genetic manipulation of livestock genomes, thereby assisting direct investigation of the genetics underpinning important phenotypes in biologically relevant cell types. We have demonstrated the utility of porcine and bovine PSCs as a source of macrophages and illustrated how they could be exploited to investigate the genetic and molecular basis of important host-pathogen interactions in livestock.

Macrophages were generated from pig and bovine PSCs using a three phase protocol, adapted from a method devised for mouse and human PSCs. Reproducibility was improved by controlling cell numbers and promoting cell association to form the embryoid body aggregates during the first phase of differentiation. The differentiation of pig PSCs typically produced ∼200 macrophages/input PSC, which means that four standard 150 cm^2^ culture flasks of PSCs could produce the ∼10^10^ macrophages equivalent to the number of alveolar macrophages harvested from a typical adult pig. PSCdMs were usually produced for 4-5 weeks, after which the cultures became exhausted. This limit to macrophage production has been observed for PSC from different species and with different differentiation protocols and might therefore reflect an intrinsic characteristic of these early haematopoietic progenitor cells^44–46^. PSC-derived myeloid progenitors and macrophages are believed to represent the *in vitro* equivalents of a transient wave of extraembryonic haematopoiesis^23^, and what determines the duration of this wave *in vivo* or *in vitro* is not yet clear. An improved understanding of the molecular mechanisms that regulate this early phase of embryonic haematopoiesis, combined with refinement of current differentiation protocols and adaptation to larger scale culture systems such as spinner cultures^45,46^, should extend and maximise the production of PSCdMs *in vitro*.

The pig and bovine PSCdMs expressed markers typical of *in vivo* macrophages, and RNA-Seq analysis indicated that the pig PSCdM transcriptional profile overlapped significantly with alveolar macrophage populations. Macrophage gene expression and phenotype is influenced by the cells ontogeny, but is also dynamic, shaped by cytokine signalling and cellular environment (e.g. substratum)^47–49^. Classically, macrophages are categorised in relation to polarisation states ranging between a M1 pro-inflammatory cell and a M2 cell involved in dampening down inflammation and promoting tissue repair^2^. However, in culture macrophages may default to a more indeterminate naive basal state and do not precisely align with a particular *in vivo* population^47^. Nonetheless, it has been reported that cultured macrophages can adopt a more *in vivo* phenotype when transplanted back into tissues *in vivo*^47^. Similarly, PSC-derived macrophages treated with the cytokines IL-34 and GM-CSF, and co-cultured with neural cells will adopt a ramified morphology and expression profile characteristic of microglial cells, the resident macrophages normally found in the brain^47,49,50^. This demonstrates that the phenotype of PSCdMs can be manipulated and exploited in culture by controlling their environment and therefore have the potential to adopt more differentiated features of tissue resident macrophages.

Notwithstanding their *in vitro* origins, livestock PSCdMs displayed many key functional attributes of *ex vivo* macrophages. The pig PSCdMs responded to immunomodulatory signals, were highly phagocytic and rapidly killed engulfed bacteria. Importantly, pig PSCdMs also served as targets for infection by key pig pathogens, including *Salmonella*, the protozoan *Toxoplasma gondii*, and the viruses ASFV and PRRSV. The obligatory PRRSV fusion receptor CD163 was expressed on PSCdMs and presumably contributed to the high levels of PRRSV infectivity achieved in these cultures. The efficient replication of ASFV and PRRSV in PSCdMs provides new opportunities to study the interactions between host genetics and the biology of these important viruses. In addition, modulation of PSCdM phenotype and infection could serve as a system for producing virus and contribute to the development of live attenuated virus vaccines and the design of novel strategies to combat diseases caused by these pathogens.

Genetic modification of pig stem cell derived macrophages was achieved by gene editing both in the undifferentiated parental stem cells and directly in PSCdMs, and provides the opportunity for functional interrogation of host genetics in a targeted manner, or through larger scale mutational screens^41,51^. This technology also affords opportunities to generate bespoke engineered cells to increase the degree of precision of experiments and for use in biotechnological applications. Although deletion of *IRF3* alone did not dramatically alter the response of PSCdMs to treatment with poly(I:C) or PRRSV infection, manipulation of the IFN response in this way could be exploited further to study the specific contribution of individual factors in mediating an antiviral or antimicrobial response. Genetically modified PSCdMs might also support enhanced replication and production of viruses, and act as more effective hosts for lentiviral-based genetic screens. Pig PSCdMs are readily transduced by lentiviral vectors, and the generation of PSCs and derivative PSCdMs that stably express Cas9 should further improve the efficiency of CRISPR/gRNA-based mutational screens^41^. The combination of bespoke genetically engineered PSCdMs and large-scale screens potentially represents a powerful approach for dissecting host-pathogen interactions.

Livestock PSCs can deliver limitless numbers of normal differentiated cells, providing a new *in vitro* platform for advancing livestock research, as well as reducing the requirement for animals as a source of primary cells or as experimental subjects. Further development of PSC-based experimental systems to produce different cell types, and their application to more complex co-culture and 3D organoid systems, affords new opportunities for functional interrogation of the molecular basis of many biologically relevant phenotypes in culture. We expect that future use of PSC-based “livestock in a dish” platforms will increase our understanding of livestock biology, and ultimately help to improve the healthy and ethical production of farmed animals.

## Materials and Methods

### Pig PSC culture

Pig PSCs were cultured on a layer of mitotically-inactivated mouse STO feeder cells (plated on gelatinised tissue culture plastic at a density of 4×10^4^/cm^2^) in pEPSC medium^27^. PSCs were passaged by washing once with PBS then incubating for 3 minutes in 0.025% trypsin/EDTA at 37°C/5%CO_2_. Cells were dispersed to single cell by pipetting and pelleted in an equal volume of feeder medium [G-MEM (Sigma, #G5154), 10% FBS (Gibco, #10500064) 1xNEAA (Gibco, #11140035), 1 mM sodium pyruvate (Gibco, #11360039), 2 mM L-glutamine (Gibco, #25030024), 0.1 nM β-mercaptoethanol (Gibco, #31350010)] at 300 x *g* for 4 minutes. Cells were plated at a density of 2-3×10^3^/cm^2^ in pEPSCM medium^27^ containing the Rho-associated coiled kinase (ROCK) inhibitor Y-27632 (5μM, Stemcell Technologies, #72304). Cells were fed the following day with pEPSCM without Y-27632 then fed daily. PSCs were passaged every 3-5 days. Two pig PSC lines (F1 and K3) were used to generate in vitro-derived macrophages^27^.

### Bovine PSC culture

Bovine PSCs (line #A)^28,31^ were cultured in bESC culture medium (bESCM) [N2B27 medium, 1xNEAA, 1xGlutamax (Gibco, #35050061) 0.1 nM β-mercaptoethanol, pen/strep (Gibco, #15140122), 10% AlbumiNZ low fatty acid BSA (MP Biochemicals, #0219989925), 20 ng/μl rhFGF2 (Peprotech, #100-18B), 20 ng/μl rhActivinA (Qkine, Qk001), 2.5 μM IWR-1 (Sigma, #I0161)] on a layer of mitotically-inactivated MEFs (plated on gelatinised tissue culture plastic at a density of 5×10^3^/cm^2^). The MEFs were washed twice with PBS prior to plating PSCs in bESCM. For passaging, PSCs were incubated 1 hour with bESCM containing 10 μM Y-27632 prior to dissociating then washed twice with PBS and incubated for 3 minutes in TrypLE Express (Gibco, #12604013) at 37°C/5%CO_2_. Cells were dispersed to single cell by pipetting and pelleted in 6x volume of bESCM at 300 x *g* for 4 minutes. Resuspended cells were plated at 1:5 in bESCM + 10 μM Y-27632 overnight then changed to bESCM without Y-27632 and fed daily. Bovine PSCs were passaged every 3-4 days.

### Macrophage differentiation

PSCs were passaged as normal then pre-plated on a gelatinised 6-well tissue culture plate for 10-15 minutes at 37°C/5%CO_2_ to remove feeder cells. Floating PSCs were pelleted at 300 x *g* for 4 minutes, washed in PBS and resuspended in StemPro (Thermo, #A1000701), 20 ng/ml rhbFGF (Qkine, #Qk027), 50 ng/ml rhBMP4 (R&D, #314-BP), 50ng/ml rhVEGF (R&D, #293-VE), 20 ng/ml rhSCF (R&D, #255-SC), pen/strep (Mesoderm Induction medium) containing 5 μM Y-27632. Typically, 2000-4000 PSCs were dispensed per well into a 96-well V-bottomed plate containing 100 μl Mesoderm Induction medium with 5 μM Y-27632 and centrifuged at 1000 x *g* for 3 minutes. The aggregated EBs were fed the next day with Mesoderm Induction medium without Y-27632 then daily thereafter. On day 4 medium was aspirated from the wells and 10-15 EBs transferred to a gelatinised 6-well tissue culture plate containing Macrophage Induction media. For porcine PSCdM differentiation, EBs were plated in medium composed of X-Vivo 15 (Lonza, #LZBE02-060F), 2 mM Glutamax, 50 nM β-mercaptoethanol, pen/strep, 100 ng/ml recombinant porcine M-CSF (Roslin Technologies), 25 ng/ml rpIL-3 (Kingfisher Biotech, #RP1298S). For bovine PSCdM differentiation, EBs were plated in medium composed of RPMI-1640 (Sigma, #R5886), 10% FBS, 2 mM Glutamax, 0.1 nM β-mercaptoethanol (cRPMI medium) containing 100 ng/ml rpM-CSF (Roslin Technologies), 25 ng/ml rpIL-3 (Kingfisher Biotech, #RP1298S). Attached EBs were fed every 4 days with Macrophage Induction medium. Early signs of macrophage production can usually be detected at day 9-12 in the form of a few attached vacuolated cells or clusters of round cells with small projections. Floating immature macrophages can typically be harvested around day 20 and collected every 4 days until approximately day 40. Harvested immature macrophages can be matured by plating cells on non-coated tissue culture plastic in X-Vivo 15, 2mM Glutamax, pen/strep, 100 ng/ml rpM-CSF (Macrophage Maturation medium).

### Phagocytosis assay

PSCdMs and PAMs were plated in triplicate on non-coated 96-well tissue culture plates at 1×10^5^/well in 100 μl Macrophage Maturation medium or RPMI-1640 (Sigma, #R5886), 10% FBS, 2 mM Glutamax, 0.1 nM β-mercaptoethanol (cRPMI medium) respectively for 48 h. On the day of the assay the medium was aspirated and replaced with 100 μl OptiMEM containing 100 μg/ml pHrodo Red Bioparticles (Thermo, #P35364). Cells were incubated at 37°C/5%CO_2_ and fluorescence measured at T0 and every hour thereafter on a BioTek Gen 5 plate reader. After 8 h the cells were dissociated by scraping with a pipette tip and fluorescence was quantified by flow cytometry. Cells with no beads added and beads alone served as negative controls.

### LPS/Poly(I:C) induction

Cells were plated at a density of 5×10^4^/cm^2^ on tissue culture plastic in Macrophage Maturation medium for 48 h. Medium was replaced with fresh Macrophage Activation medium containing either Lipopolysaccharides (LPS, 200 ng/ml, from *Escherichia coli* O111:B4, Sigma #L4391) or Poly(I:C) (25 μg/ml, Tocris #4287) and incubated for 4 h prior to lysis for RNA recovery.

### RT-qPCR

RNA was prepared using Qiagen RNeasy kit (#74104) following the manufacturer’s protocol including the recommended on-column DNase treatment. cDNA was synthesised from 0.2-1 μg of RNA using Agilent’s Multitemp cDNA Synthesis kit (#200436) at 42°C following the manufacturer’s instructions. The final cDNA volume was made up to 1 ml with nuclease-free water. Each RT-qPCR reaction consisted of 8 μl of diluted cDNA plus a mastermix consisting of 10 μl Agilent Brilliant III SYBR green (#600883), 0.4 μl Reference dye (2 μM) and 0.8 μl each of forward and reverse primers (RPL4 was used as the housekeeping gene to normalise expression - see list of primers). The reaction was performed on a Stratagene MxPro3005P QPCR instrument using the following cycle parameters: one cycle of 95°C for 2 minutes followed by 40 cycles of 95°C for 15 s and 60°C for 30 s. A final cycle of 95°C for 1 minute, 60°C for 30 s and 95°C for 15 s was performed to establish a dissociation curve.

### Cell surface staining

Cells were blocked in PBS/2%FBS, on ice for 30 minutes. 2×10^5^ cells/well were transferred to a 96-well V-bottomed plate and pelleted at 300 x *g*, 4 minutes, 4°C. The supernatant was removed by inverting the plate. The pellet was resuspended with 25 μl of diluted, conjugated antibody and incubated in the dark, on ice for 30 minutes. The cells were then pelleted at 300 x *g*, 4 minutes, 4°C and washed twice with 75 μl PBS. The final pellet was resuspended in 100 μl PBS. 100μl SYTOX Blue Nucleic Acid Stain (5μM, ThermoFisher #S11348) was added immediately prior to flow cytometer analysis to allow for live/dead cell identification. Antibodies used were CD14 (Biorad, #MCA1218F, 1:50) with isotype control (Sigma, #SAB4700700), CD16 (Biorad, #MCA1971PE, 1:200) with isotype control (Biorad, #MCA928PE), CD163 (Biorad, #MCA2311F, 1:100) with isotype control (Sigma, #F6397), CD169 (Biorad, #MCA2316F, 1:100) with isotype control (Sigma, #F6397) and CD172a (Southern Biotech, #4525-09, 1:400) with isotype control (Biorad, #MCA928PE).

### *Toxoplasma* infection and staining

Pig PAMs and PSCdMs were plated 48 h prior to infection at 8×10^5^/well in a 12-well tissue culture plate in cRPMI medium. The cells were fed with cRPMI 24 h before infection. The next day the medium was aspirated and the cells infected with *Toxoplasma gondii* at MOI=1 in cRPMI for 24 h at 37°C/5%CO_2_. 24 h post-infection the cells were collected using a cell scraper and pelleted at 600 x *g* for 4 minutes then washed in 500 μl PBS. One half was lysed for genomic DNA recovery to determine *Toxoplasma* DNA copies and the other half used to prepare RNA for RT-qPCR analysis.

### *Salmonella enterica* serovar Typhimurium infection and staining

Pig PAMs and PSCdMs were plated 48 h prior to infection at 5×10^5^/well in a 12-well tissue culture plate in cRPMI. The day before infection, a single colony of *Salmonella enterica* serovar Typhimurium strain 4/47, expressing EGFP from plasmid pFVP25.1^32^, was cultured for 16 h in 3 ml LB medium + 100 μg/ml Ampicillin. The OD_600_ absorbance was measured on a spectrophotometer and used to determine the bacterial cell concentration using the online tool http://www.labtools.us/bacterial-cell-number-od600/. The cells were washed twice with PBS and infected with bacteria diluted in cRPMI medium at an MOI=2 for 30 minutes at 37°C/5%CO_2_. Following two washes with PBS the cells were then treated with 100 μg/ml gentamicin in cRPMI for 1 h at 37°C/5%CO_2_ to kill extracellular bacteria. Surviving intracellular bacteria were harvested at 0 h and 3 h after gentamicin treatment by washing the cells twice with PBS then lysing with 1% TritonX100. 10-fold serial dilutions were plated on to LB/Ampicillin culture plates and incubated overnight at 37°C. Colonies were counted the next day. For staining, cells were fixed on glass coverslips after gentamicin treatment in 4% Formaldehyde for 15 minutes then permeabilised in PBS/0.1% TritonX100 for 10 minutes before staining with Phalloidin AF647 (1:1000) and DAPI (1:10,000) in PBS at room temperature for 30 minutes in the dark. Cells were washed twice with PBS before mounting on glass slides and imaging on a Leica LSM10 confocal microscope.

### *Escherichia coli* infection

Pig PAMs and PSCdMs were infected with *Escherichia coli* strain TOP10 at a MOI=10 using the same protocol as for *Salmonella* infection and surviving bacteria harvested at 0 h and 2 h after gentamicin treatment.

### PRRSV infection and staining

Pig PAMs and PSCdMs were plated on non-coated tissue culture plates in cRPMI at a density of 1×10^5^/cm^2^ 24 h prior to infection. Cells were infected with PRRSV (SU1-Bel) at MOI=1 in cRPMI for 2 h at 37°C. The inoculum was then removed, and the cells fed with fresh cRPMI. At 19 hpi cells were washed twice with PBS and detached using a cell scraper. Cells were fixed in 4% Formaldehyde for 15 minutes then permeabilised with 0.1% TritonX100/PBS for 10 minutes. After washing twice with PBS, the cells were blocked in 5%FBS/PBS for 30 minutes prior to incubating with primary antibody (SDOW-17A, 1:5000) for 45 minutes in blocking solution. Following two washes with PBS, the cells were then incubated with secondary antibody (Goat α-mouse AF488, 1:5000) for 1 h in the dark before staining with Phalloidin AF647 (1:1000) and DAPI (1:10,000) for 30 minutes in the dark. After two washes with PBS the cells were analysed by flow cytometry.

### ASFV infection and growth assays

Porcine monocyte macrophages (PMMs) were harvested from heparinised blood taken from pigs housed at the APHA under housing and sampling regulations, licence PP1962684, approved by the APHA Animal Welfare and Ethical Review Board and conducted in accordance with the Animals (Scientific procedures) Act UK. Blood was centrifuged and plasma, leukocytes (buffy coat) and erythrocyte fractions harvested. The leukocytes were washed in PBS, followed by two washes with BD Pharm Lyse (#555899). After two further washes in PBS cells were re-suspended in RPMI supplemented with 20% v/v autologous plasma, harvested from the initial centrifugation step, and 100U/ml Penicillin-Streptomycin (Gibco, #15140122). Cells were incubated in 96-well plates at 37 °C/5% CO_2_ for 48h prior to infection with ASFV. Infections with ASFV were performed at the APHA in biosecure containment laboratories licenced for handling of level 4 specified animal pathogens. ASFV strain Armenia 07 diluted in RPMI was added to PSCdMs, PAMs and PMMs at an MOI of 1 in 96-well plates. After 1 hour incubation at 37°C the virus inoculum was removed and, for quantification of viral replication by qPCR, was replaced with 200μl of Macrophage Maturation media. For observation of ASFV infection by detection of haemadsorbance additional wells were set up in which the virus inoculum was removed and replaced with 200μl Macrophage Maturation medium supplemented with 1% v/v porcine erythrocytes and 1% porcine plasma. Plates were incubated at 37°C/5% CO_2_ for up to 5 days. Formation of HAD rosettes due to haemadsorbance of erythrocytes to infected macrophages was observed by light microscopy. To quantify ASFV replication and release into the supernatant 140μl of media was removed from wells after 0, 24 and 48 hours and nucleic acid extracted using Qiamp viral RNA mini extraction kit (Qiagen, #52904). Viral DNA levels were quantified by qPCR using primers and probe that detect the ASFV VP72 gene^52^ with the Quantifast Pathogen PCR kit (Qiagen) and the following cycle conditions: 1x 95°C for 5min followed by 50 cycles of 95°C for 15 s, 60°C for 1min. The copies of viral genome were determined by comparison Cq values to those of a standard comprised of a dilution series of the plasmid pASFV-VP72 encoding a fragment of the VP72 gene.

ASFV infection with strain Benin 97/1 was performed at the Pirbright Institute essentially as described previously^53^. ASFV infection was monitored by formation of HAD rosettes, and immunocytochemical detection of ASFV VP72 expression^54^. ASFV replication was measured by TCID_50_ assay and calculated using the Spearman-Karber method.

### Gene editing *REX1-EGFP knock-in*

The pig *REX1* targeting vector was constructed in two stages. First, the homology arms were amplified from pig PSC genomic DNA using primers with tails containing the inverted guide sequence (5’HA+Guide_Forward:CTTCTTTCACTGATTTGTATTGGTTCAAGGAGAGCGCAAAACTA,3’HA+guide_Reverse:CTTCTTTCACTGATTTGTATTGGAGTTGATTCAAATGGATTGACA). The PCR product was then TA-cloned into the pCR4-TOPO TA vector backbone (ThermoFisher #450071) and linearised by inverse PCR using primers positioned either side of, and designed to exclude, the Rex1 STOP codon (HA3_inv_For AAGAAGACTGAAAATAATCC, HA3_inv_Reverse:CTGATTTGTATTGGCCTTTG). In addition, a T2A-EGFP-IRES-PURO-bGHpA cassette was amplified by PCR using primers with 15 bp tails homologous to the sequence either side of the Rex1 STOP codon (T2ARex1_Forward:GCGAATACAAATCAGGGCTCCGGAGAGGGCAGAG, bGHpaRex1_Reverse:ATTTTCAGTCTTCTTCCATAGAGCCCACCGCATCC). Second, the linearised homology arms and amplified reporter/selection cassette were assembled by Gibson assembly (NEB, #E2621S) and individual clones were sequence verified. A CRISPR/Cas9 guide sequence was identified using Benchling (www.benchling.com) that generates a double-strand break 8 bp upstream of the pig *REX1* STOP codon (Rex1_363 CTTCTTTCACTGATTTGTAT). The sgRNA was synthesised by Synthego. For editing, 7.5 μl sgRNA (100 μM) was combined with 5 μl Cas9 protein (20 μM, Synthego) at room temperature for 10 minutes to form ribonucleoprotein complexes (RNPs) then 1 μg targeting vector was added to the RNPs and made up to 30 μl with P3 Primary Cell Solution (82 μl Nucleofector Solution + 18 μl Supplement per 100 μl) and kept on ice prior to transfection. Pig PSCs were passaged as normal and 5×10^5^ cells were resuspended in 70 μl of Amaxa P3 Primary Cell Solution. The RNP complex was mixed with the cells, transferred to a transfection cuvette then nucleofected on an Amaxa 4D Nucleofector using program CG-104. The cells were resuspended in pEPSCM + ROCKi and plated over two wells of a 6-well plate containing mitotically-inactivated STO feeder cells. Medium was changed the next day for pEPSCM without ROCKi. 72 h post-transfection the cells were passaged and plated at 2×10^4^/cm^2^. Puromycin selection (0.2 μg/ml) was added 24 h later. After 10 days six colonies were picked and passaged as normal into a 96-well tissue culture plate. Clones were expanded and screened by PCR for evidence of editing. Correctly targeted clones were identified at both the 5’ an 3’ ends of the integration site by PCR amplification of genomic DNA using 5’ primers (xF1 - GTTTTCTGAGTACGTGCCAGGC, iR1 - CGGGTCTTGTAGTTGCCGTCGT) and 3’ primers (iF2 - TGGGAAGACAATAGCAGGCATG, xR3 – CACACCCCGCCCAACTGCTG) under the following cycle conditions - 98°C for 1 minute then 32 cycles of 98°C for 10 s, 69°C for 30 s and 72°C for 1 minute followed by a final extension of 72°C for 10 minutes. For each screen one primer was located outside the homology arm sequence and the other within the reporter/selection cassette. Targeted fragments of 885 bp and 1213 bp were expected for the 5’ and 3’ screens respectively.

### Gene editing *IRF3* deletion

A pair of CRISPR/Cas9 guide sequences were designed to delete the pig *IRF3* coding sequence. Guide sequences were identified using Benchling (www.benchling.com) and synthesised by Synthego (IRF3_1094 CGAGGCTTCTGAGTTCCCAT, IRF3_5441 ACATGGATTTCTAGGCCGCT). For editing, 3.75 μl of each sgRNA (100 mM) was combined with 5 μl Cas9 protein (20mM, Synthego) at room temperature for 10 minutes to form RNPs then 17.5 μl P3 Primary Cell Solution added and the RNPs kept on ice prior to transfection. Pig PSCs were passaged as normal and 5×10^5^ cells were resuspended in 70 μl of Amaxa P3 Primary Cell Solution. The RNP complex was mixed with the cells, transferred to a transfection cuvette then nucleofected on an Amaxa 4D Nucleofector using program CG-104. The cells were resuspended in pEPSCM + ROCKi and plated over two wells of a 6-well plate containing mitotically-inactivated STO feeder cells. Medium was changed the next day for pEPSCM without ROCKi. 72 h later the cells were passaged and plated at low density (2.5×10^2^-1×10^3/^cm^2^). After 9-11 days 80 colonies were picked and passaged as normal into a 96-well tissue culture plate. Clones were expanded and screened by PCR for evidence of editing. Genomic DNA was PCR amplified using a pool of two forward primers (scrnF1 - AGGCCGTCTGTTTGGGAGGAA, Ex8F1 - TTGTCCCCATGTGTCTCCGG) and one reverse primer (scrnR1 - TGACAGACAGGACGTTTAGGCA) under the following cycle conditions - 98°C for 1 minute then 32 cycles of 98°C for 10 s, 68°C for 30 s and 72°C for 1 minute followed by a final extension of 72°C for 10 minutes. Two wild-type fragments of 5,311 bp and 640 bp, and an edited fragment of 964 bp were expected although the 5,311 bp fragment failed to amplify under these conditions.

### Lentivirus Packaging

HEK293T cells were grown to 80% confluence in a T175 flask then transfected with 15 μg lentiviral plasmid together with 12 μg psPax2 and 3 μg pVSV packaging plasmids using 15 μl Lipofectamine 2000. The medium containing lentivirus was harvested at 24 h and 48 h. The medium was stored at 4°C until all harvests were collected then pooled and filtered through a 0.45 μm filter. Filtered lentivirus was either stored in aliquots at −80°C or further purified and concentrated using the Lenti-X Maxi Purification kit (Takara #631234) according to the manufacturer’s instructions. Concentrated lentivirus was stored in aliquots at −80°C.

### Lentiviral transduction

Pig PSCdMs were plated in Macrophage Maturation Medium at 3×10^5^/cm^2^. 72 h later, the medium was removed, and the cells were transduced with lentivirus (250 μl/cm^2^) in Macrophage Maturation Medium containing 2 μg/ml Polybrene (Santa Cruz, sc134220) by spinfection (centrifugation at 1000 x *g* for 1 h at 32°C). Following spinfection, the medium was replaced with fresh Maturation Medium, and the cells incubated at 37°C/5%CO_2_. The cells were imaged and analysed by flow cytometry 7-8 days post-transduction. For assessing transduction efficiency a CMV-GFP-Puro-expressing lentivirus (Addgene #17448) was used at MOI=1. For editing of CD163 a dual guide RNA lentivirus (Addgene #67974) was modified to express the CD163 guides SL26 and SL28^55^ by cloning a gBlock containing the crRNASL26-tracrRNA-mU6-crRNASL28 sequence into the *Bbs*I site. The CD163 guide lentivirus was co-transduced along with the Cas9-expressing lentivirus, lenti-Cas9-Blast (Addgene #52962) at 1:1 v/v. The empty dual guide lentivirus was used as a negative control. CRISPR/Cas9-mediated deletion of CD163 exon7 was determined by genomic DNA PCR amplification using primers (CD163scrnF - ACCTTGATGATTGTACTCTT, CD163scrnR - TGTCCCAGTGAGAGTTGCAG) under the following cycle conditions 98°C for 1 minute then 32 cycles of 98°C for 10 s, 67°C for 30 s and 72°C for 1 minute followed by a final extension of 72°C for 10 minutes. A wild-type fragment of 941 bp, and an edited fragment of 454 bp were expected.

### Bioinformatics

Pig RNA-seq datasets used for estimating gene expression were obtained from NCBI (BioProject: PRJEB19386 and GEO: GSE172284^56^). Illumina short-read RNA-Seq data was adapter trimmed^57^ and aligned to the pig reference genome (Sscrofa11.1^58^) using STAR (v 2.7.1a)^59^ only allowing a maximum of 20 multimappers per read. Mapping rates were consistently above 90%. The number of mapped reads were counted at gene level using featureCount (v. 1.6.3)^60^ with the Ensembl pig genome annotation (v.101)^61^. Heat map, sample specific clustering and PCA plots were created in R (https://www.R-project.org/.) using the DESeq2 package^60^. Genes of low or now expression were filtered out (total read counts per gene < 20) and a variance stabilizing transformation was used before comparing the samples.

### Pig RT-qPCR primer list

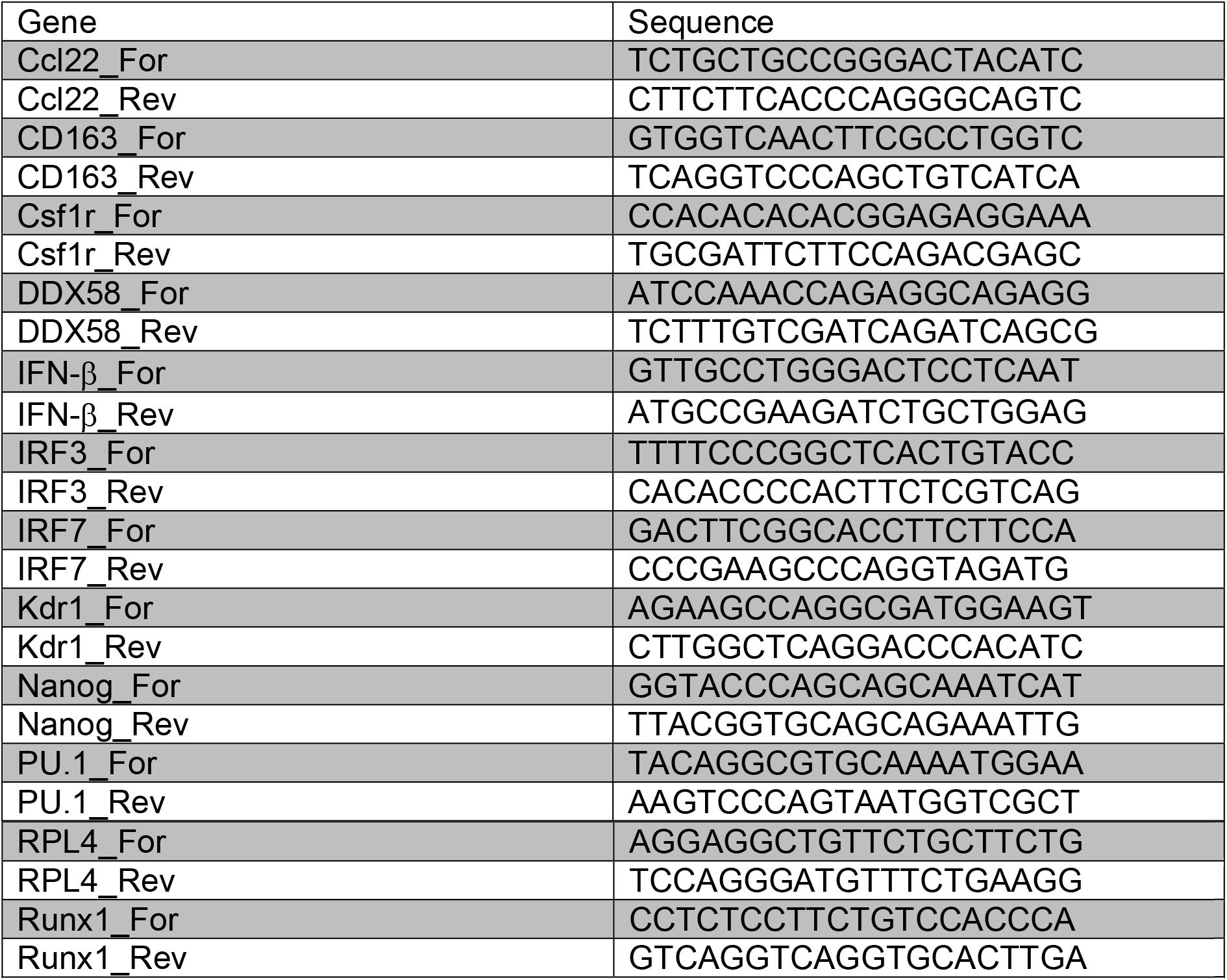

### Bovine RT-qPCR primer list

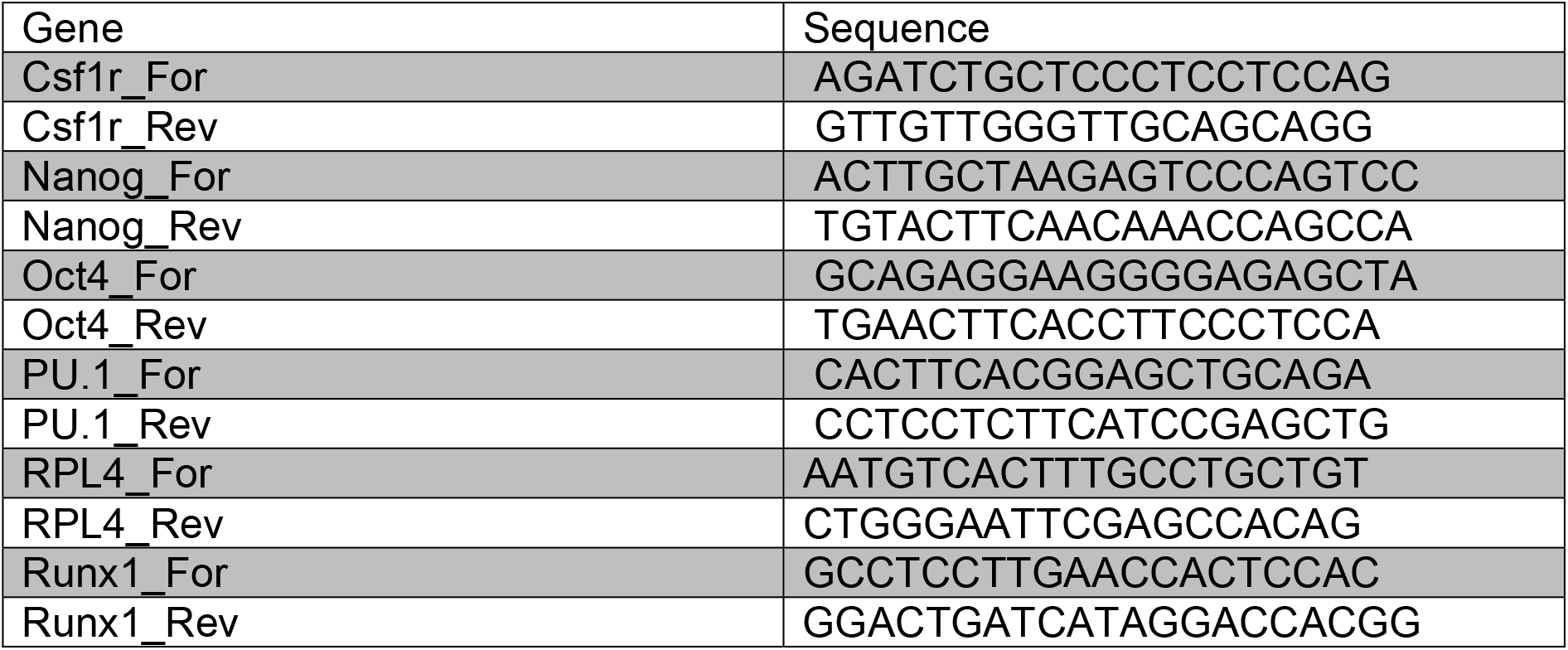

## Acknowledgments

This work was supported by funding from the Biotechnology and Biological Sciences Research Council Institute Strategic Programme Grant BB/P013732/1; a BBSRC Industrial CASE EASTBIO PhD studentship; BBSRC Impact Accelerator Award to the University of Edinburgh PIII054; and National Centre for the Replacement, Refinement and Reduction of Animals in Research (NC3Rs) grant NC/V001140/.

## Author contributions

Stephen Meek: Conceptualization; Data curation, Formal analysis, Validation, Investigation, Methodology, Supervision, Writing—review and editing; Tom Watson: Investigation, Methodology; Lel Eory: Methodology, Data curation, Formal Analysis; Gus McFarlane: Investigation; Felicity J Wynne: Investigation, Formal analysis; Stephen McCleary: Investigation; Laura E.M. Dunn: Investigation, Formal analysis; Emily M. Charlton: Investigation; Chloe Craig: Investigation; Barbara Shih: Data curation, Formal Analysis; Tim Regan: Methodology; Ryan Taylor; Methodology; Linda Sutherland: Methodology; Anton Gossner: Methodology, Resources; Cosmin Chintoan-Uta: Methodology, Resources; Sarah Fletcher: Methodology, Resources; Philippa M. Beard: Methodology, Resources, Supervision; Musa A. Hussan; Methodology, Resources, Supervision; Finn Grey: Methodology, Resources; Jayne C. Hope; Methodology, Supervision; Mark P Stevens: Methodology, Resources; Monika Nowak-Imialek: Resources; Heiner Niemann; Resources; Pablo J. Ross: Resources; Christine Tait-Burkard: Methodology, Resources, Supervision; Sarah M. Brown: Resources; Lucas Lefevre: Resources; Gerard Thomson: Resources; Barry W McColl: Resources; Alistair B Lawrence: Resources; Alan L. Archibald: Resources, Supervision, Review and editing; Falko Steinbach: Resources, Supervision; Helen R. Crooke: Methodology, Resources, Supervision, Formal analysis; Xuefei Gao: Resources, Methodology; Pentao Liu: Resources, Supervision; Tom Burdon: Conceptualization, Funding acquisition, Project administration, Supervision, Data curation Formal analysis, Writing original draft, Writing—review and editing.

---

**Supplementary Figure 1.**
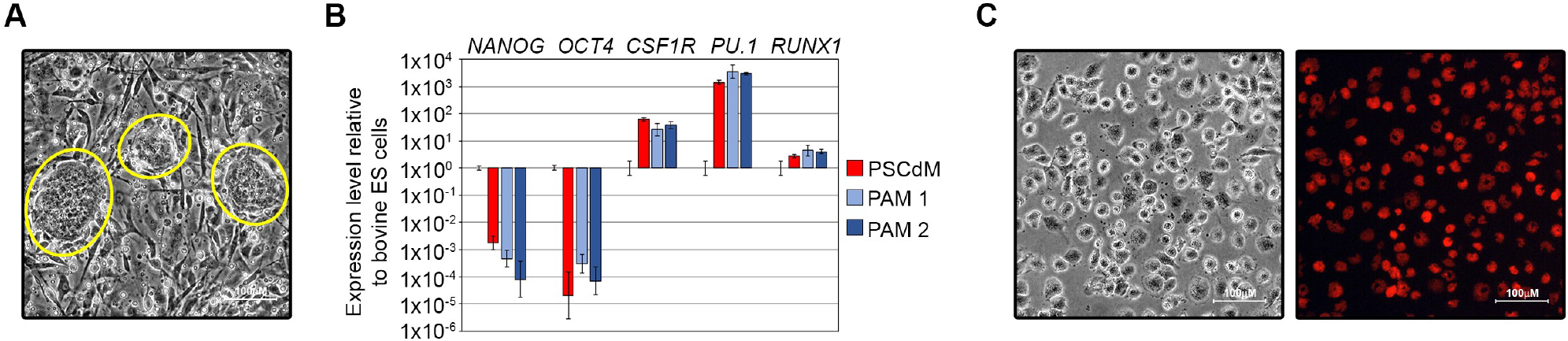
**(A)** Bright-field image of bovine PSCs grown on mitotically-inactivated MEFs. Bovine PSC colonies are circled in yellow. **(B)** RT-qPCR analysis comparing expression of pluripotency markers (*NANOG* and *OCT4*) and macrophage markers (*CSR1R, PU*.*1* and *RUNX1*) in primary bovine PAMs^62^ and bovine PSCdMs relative to bovine PSCs. Mean and SD of three technical replicates. **(C)** Bright-field and fluorescent images of bovine PSCdMs containing phagocytosed pHrodo beads.

**Supplementary Figure 2 (related to figure 2F).**
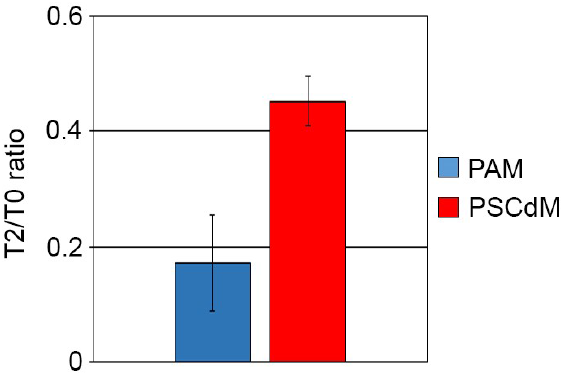
Ratio of colony-forming *Eschericia coli* recovered from infected primary pig PAMs and pig PSCdMs at 2 hr post-infection relative to T0. Mean and SD of duplicate plates from two experiments.

**Supplementary Figure 3 (related to figure 3C & D).**
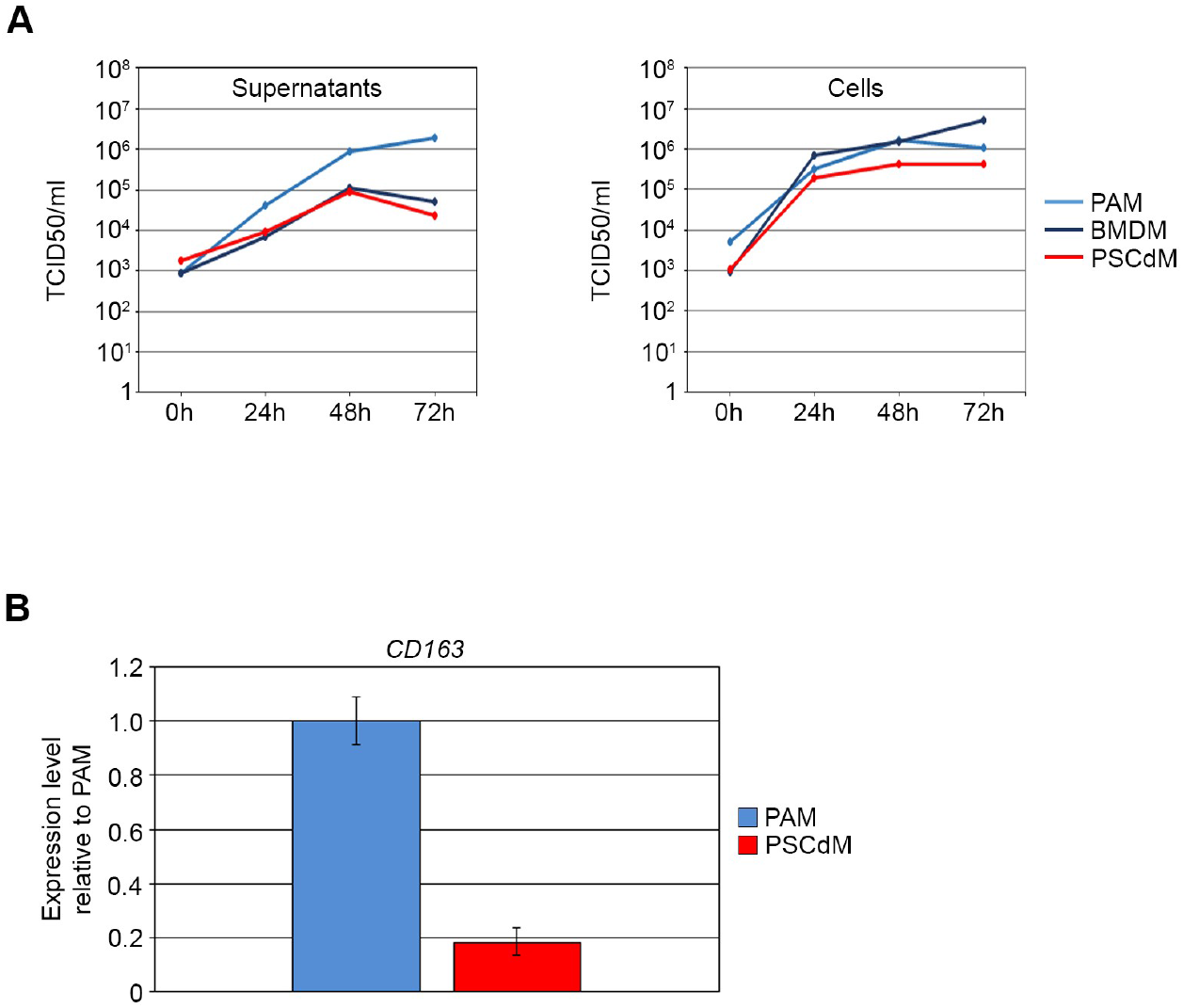
**(A)** Pig PAMs, BMDMs and PSCdMs were infected with ASFV (Benin 97/1 strain). Viral replication was determined by harvesting both supernatants and cells at 0, 24, 48 and 72 hpi, and titrating on pig BMDMs. TCID_50_ was calculated by the Spearman-Karber method. Data points represent mean of experimental duplicates. **(B)** RT-qPCR analysis comparing *CD163* expression in primary pig PAMs and pig PSCdMs. Mean and SD of duplicate samples from two experiments.

**Supplementary Figure 4.**
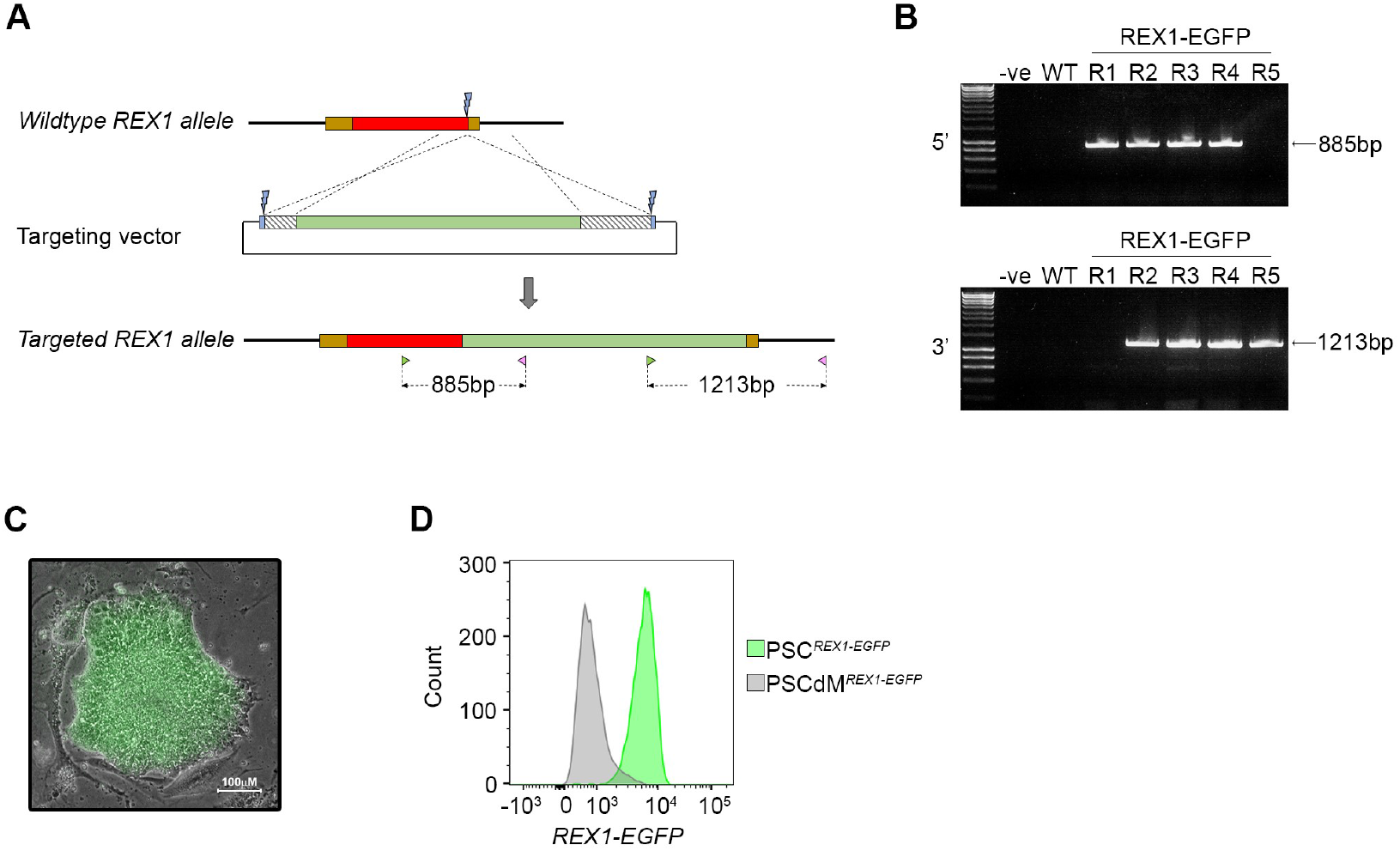
**(A)** Targeting diagram showing wild-type (top) and targeted (bottom) pig *REX1* alleles generated using the PITCh targeting vector (middle) following CRISPR/Cas9-mediated homology-directed repair as indicated by the dotted lines. The targeting vector consisted of a T2A-EGFP-IRES-PURO-bGHpA cassette (green box) flanked by a 243 bp 5’ homology arm and a 534 bp 3’ homology arm (grey hashed boxes). The homology arms were flanked by inverted CRISPR/Cas9 guide sequences (blue boxes) that matched the endogenous CRIPSR/Cas9 cut site sequence (blue lightning bolts). Following co-electroporation of the targeting vector and Cas9/sgRNA RNP, puro-resistant PSC colonies were generated in which the *REX1* stop codon had been replaced with the reporter/selection cassette at the 3’ end of the *REX1* coding exon (red box) immediately upstream of the 3’ UTR (brown box). Non-coding genomic sequence and plasmid backbone sequence are represented by thick and thin black lines respectively, and 5’ and 3’ UTRs by brown boxes. Confirmation of correctly targeted clones was performed at both the 5’ and 3’ end of the integration site using forward and reverse primers flanking the 5’ and 3’ homology arms respectively. Expected PCR product sizes are indicated. **(B)** Five puro-resistant, EGFP+ clones were genotyped by PCR using the primers indicated in panel A. Clones R2, R3 & R4 showed the expected products at both the 5’ and 3’ ends of the integration site. Water and wildtype, parental pig PSC genomic DNA were used as negative controls (-ve and WT respectively). **(C)** Compound bright-field and fluorescent image of a *REX1-EGFP* positive pig PSC colony. **(D)** Flow cytometry analysis of *REX1-EGFP* pig PSCs and *REX1-EGFP* pig PSCdMs.

**Supplementary Figure 5 (related to figure 4E).**
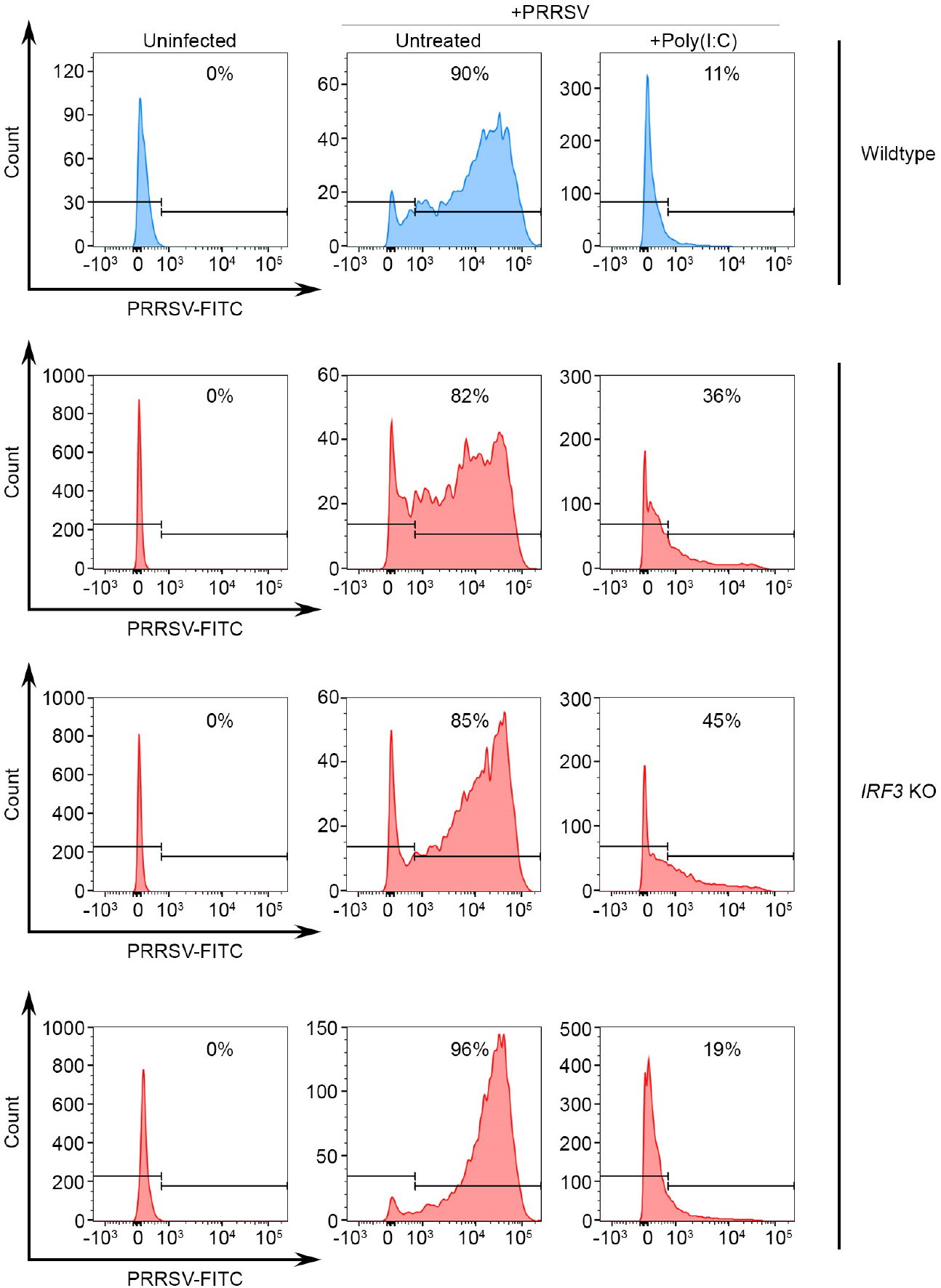
Flow cytometry analysis for PRRSV nucleocapsid protein in three *IRF3* knock-out (KO) pig PSCdMs clones relative to the wild-type parental line. Plots represent uninfected (left), untreated/infected (middle) and poly(I:C)-treated/infected (right). For poly(I:C) treatment cells were pre-treated with 25 μg/ml for 3 h prior to infection.

**Supplementary Figure 6 (related to figure 4I).**
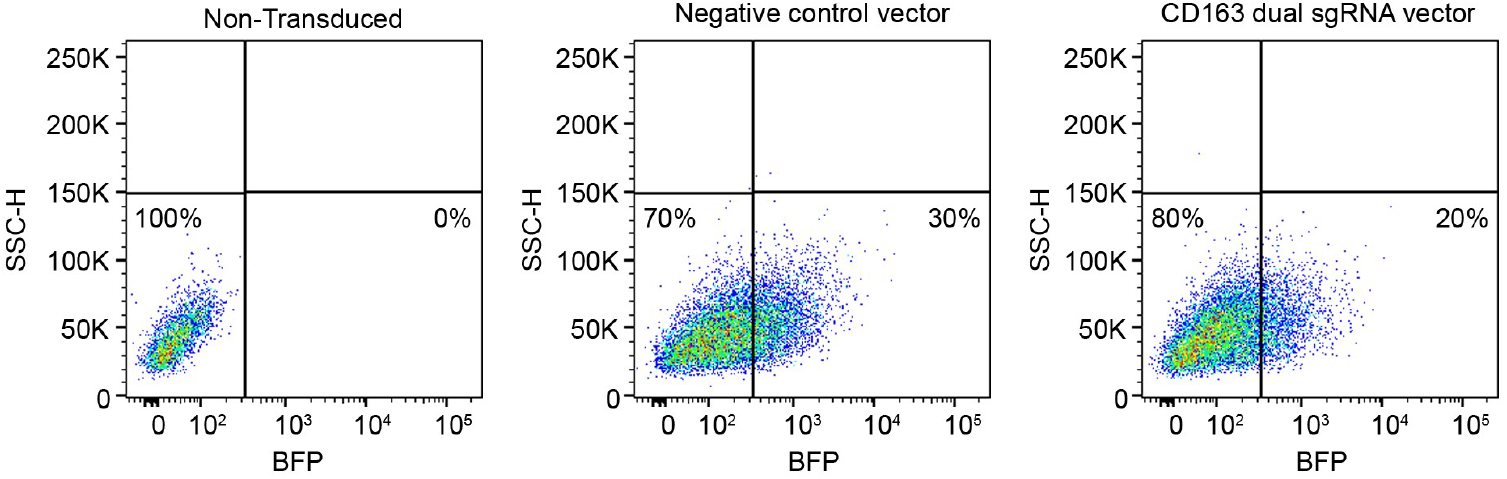
Flow cytometry data for pig PSCdMs transduced with a lentiviral dual-expression vector expressing the CD163 CRISPR guide RNAs SL26 and SL68^55^ (right panel) or a negative control vector containing no guide sequences (middle panel) relative to non-transduced cells (left panel). BFP^+ve^ cells were sorted seven days post-transduction using the conservative FACS gate shown.

